# A novel *bla*_SIM-1_-carrying megaplasmid pSIM-1-BJ01 isolated from clinical *Klebsiella pneumonia*

**DOI:** 10.1101/519082

**Authors:** Yang Lü, Hui Liang, Wei Zhang, Jia Liu, Shiwei Wang, Hongyan Hu

## Abstract

A rare carbapenem-resistant gene *bla*_SIM-1_ was found in a 316-kb megaplasmid designated pSIM-1-BJ01 isolated from a clinical strain *Klebsiella pneumonia* 13624. The plasmid pSIM-1-BJ01 was fully sequenced and analyzed. Its length is 316,557 bp and it has 342 putative open reading frames with two multidrug-resistant regions and a total of 19 resistant genes. Its backbone was highly homologous to the newly reported plasmid pRJA166a, which was isolated from a clinical third-generation cephalosporin-resistant hypervirulen strain K. pneumonia ST23. The plasmid pSIM-1-BJ01 was verified to be able to transfer to *Escherichia coli*. The emergency of the transferable *bla*_SIM-1_-carrying multidrug-resistant plasmid pSIM-1-BJ01 suggests the spread of *bla*_SIM_ among Enterobacteriaceae is possible. Therefore, the data presented herein provided insights into the genomic diversity and evolution of *bla*_SIM_-carrying plasmids, as well as the dissemination and epidemiology of *bla*_SIM_ among Enterobacteriaceae in public health system.

## INTRODUCTION

SIM, a rare member of metallo-β-lactamases (MBLs), belongs to Class B1 MBL_S_. It can hydrolyze a broad array of β-lactams, including penicillin, narrow-to expanded-spectrum of cephalosporins, and carbapenems, but not monobactams. The SIM-1 protein exhibits 64-69% identity with the IMP-type MBLs, which are its closest relatives. The first reported bacteria carrying *bla*_SIM-1_ were *Acinetobacter baumannii* isolated from Korea in 2005 (1). In *bla*_SIM-1_-harboring *Acinetobacter* isolates collected from 2003 to 2008 in Korea, most of the *bla*_SIM-1_ genes were carried in ca. 280-kb plasmids; further study found that the plasmids were transferable, but not promiscuous (2). In 2011, an *A. baylyi* strain with a 360-kb plasmid carrying both *bla*_SIM-1_ and *bla*_OXA-23_ was isolated from a Chinese patient in Ningbo of China, who had no history of visiting Korea, but the sequence of the 360-kb plasmid was unclear (3). In 2012, a *Pseudomonas aeruginosa* strain with a 282-kb *bla*_SIM-2_-harbouring plasmid pHN39-SIM (Genebank Acession Number KU254577) was isolated from a Chinese patient in Zhengzhou of China; the 282-kb *bla*_SIM-2_-harbouring plasmid was sequenced, showing the SIM-2 protein differs from SIM-1 due to only a single amino acid substitution Gly198Asp (4). Overall, the detection of *bla*_SIM_ is rare and mainly reported by South Korea and China, mostly confined to *Acinetobacter spp*. and *P. aeruginosa*.

The reported *bla*_SIM-1_ genes, derived from chromosome or plasmids, were always carried on a gene cassette inserted into a class I integron with three additional resistant genes (*arr-3, catB3* and *aadA1*). Except for the *bla*_SIM-2_-harbouring plasmid pHN39-SIM from *P. aeruginosa*, none of these plasmids is fully sequenced.

Here, we present a 316-kb plasmid designated pSIM-1-BJ01 carrying *bla*_SIM-1_ isolated from a clinical carbapenem-resistant *Klebsiella pneumonia* 13624 strain harboring three resistant plasmids in Beijing of China. To the best of our knowledge, this is the first sequence report of a transferable plasmid with *bla*_SIM-1_ isolated from *K. pneumonia*, indicating the active occurrence of resistant gene convergence under clinical selection pressure and presenting a considerable challenge for infection control.

## MATERIALS AND METHODS

### Bacterial Isolates and Identification

The isolate was obtained from a urinary specimen of a patient with 65~70 years, who suffered from the insulin-dependent type II diabetes, end-stage renal failure and pulmonary infection. The bacterial species identification was performed using BioMérieux VITEK2, Bruker MALDI Biotyper and 16S rRNA gene sequencing.

### Antimicrobial Resistance

Antimicrobial susceptibility testing was performed by VITEK-2 (bioMérieux) and evaluated according to 2014 CLSI guidelines. Metallo-β-lactamase activity was detected by Modified Hodge test. The major acquired carbapenemase genes (*bla*_GES_, *bla*_KPC_, *bla*_SME_, *bla*_IMI_, *bla*_BIC_, *bla*_IMP_, *bla*_VIM_, *bla*_NDM_, *bla*_TMB_, *bla*_FIM_, *bla*_SPM_, *bla*_DIM_, *bla*_GIM_, *bla*_SIM_, *bla*_AIM_, *bla*_SMB_, *bla*_OXA_) were screened for by PCR with specific primers described by Chen et al (5).

### Plasmid Transfer

Conjugal transfer experiment was performed with *Escherichia coli* J53 Azi^r^ as the recipient strain. Overnight cultures of the bacteria was diluted to 1.5 × 10^8^ cells/mL. Donor and recipient cells were mixed at 1:10 donor-to-recipient ratio, after 18 h of incubation of donor-recipient mixtures on blood plates at 35°C, cells were washed by normal saline solution and diluted to be cultured selectively on MacConkey agar plates supplemented with sodium azide (100 μg/mL) and imipenem (1 μg/mL) for 24 h. Transconjugants were confirmed by PCR amplifying *bla*_SIM_ with primers (SIM-F: 5’ TACAAGGGATTCGGCATCG 3’; SIM-R: 5’ TAATGGCCTGTTCCCATGTG 3’).

### Whole Genome Sequencing and Data Analysis

Bacterial genomic DNA was sequenced with Agilent 2100 and PacBio RSII platform. The contigs were assembled by SMRT Portal program. The genes were predicted with GeneMarkS and further annotated by BLASTP and BlASTN against UniPort and NR databases. The complete sequence of pSIM-1-BJ01, pSIM-1-BJ02, and pSIM-1-BJ03 was submitted to GenBank under accession number MH681289, MK158080, and MK158081, respectively.

## RESULTS

### Antibiotic Resistance of the *bla*_SIM-1_-Carrying *K. pneumoniae*

The isolated *K. pneumoniae* 13624 strain was resistant to ampicillin, cephalosporin, and carbapenem, but sensitive to ciprofloxacin, levofloxacin, and colistin (Table 1). We detected the major carbapenemase genes by PCR, only the *bla*_SIM-1_ gene was amplified.

**TABLE 1.** Antimicrobial drug susceptibility profiles.

| Antibiotic | MIC (mg/L) |  |
| --- | --- | --- |
|  | *K. pneumoniae* 13624 | *E. coli* J53 (pSIM-1-BJ01) |
| Ampicillin | >256 | >256 |
| Piperacillin | >128 | >128 |
| Cephalothin | >256 | >256 |
| Cefoxitin | >256 | >256 |
| Cefotaxime | >256 | >256 |
| Cefuroxime | >256 | >256 |
| Ceftazidime | >256 | >256 |
| Aztreonam | >128 | >128 |
| Cefepime | >64 | >32 |
| Ertapenem | >16 | >8 |
| Imipenem | >16 | >8 |
| Meropenem | >16 | >8 |
| Ciprofloxacin | <0.5 | <0.5 |
| levofloxacin | <0.5 | <0.5 |
| Colistin | <0.5 | <0.5 |

In the conjugation experiment, we also detected the *bla*_SIM-1_ by PCR in the recipient *E.coli* J53 Azi^r^, suggesting that a transferable plasmid (designated pSIM-1-BJ01) might exist in the clinically isolated *K. pneumoniae*. The transferable plasmid pSIM-1-BJ01 might give the recipient *E.coli* J53 Azi^r^ the carbapenem resistance, as shown in Table 1.

We sequenced the whole genome of the isolated *K. pneumonia* and found that there were three drug-resistant plasmids in the isolated *K. pneumonia*. They are respectively the *bla*_SIM-1_-harboring megaplasmid pSIM-1-BJ01 (Fig. 1), pSIM-1-BJ02 (Fig. 3) carrying the extended-spectum β-lactamase (ESBL) genes *bla*_TEM-1_ and *bla*_CTX-M-15_, and pSIM-1-BJ03 (Fig. 4) carrying the ESBL genes *bla*_CTX-M-14_ and *bla*_LAP-2_.

**Figure 1.**
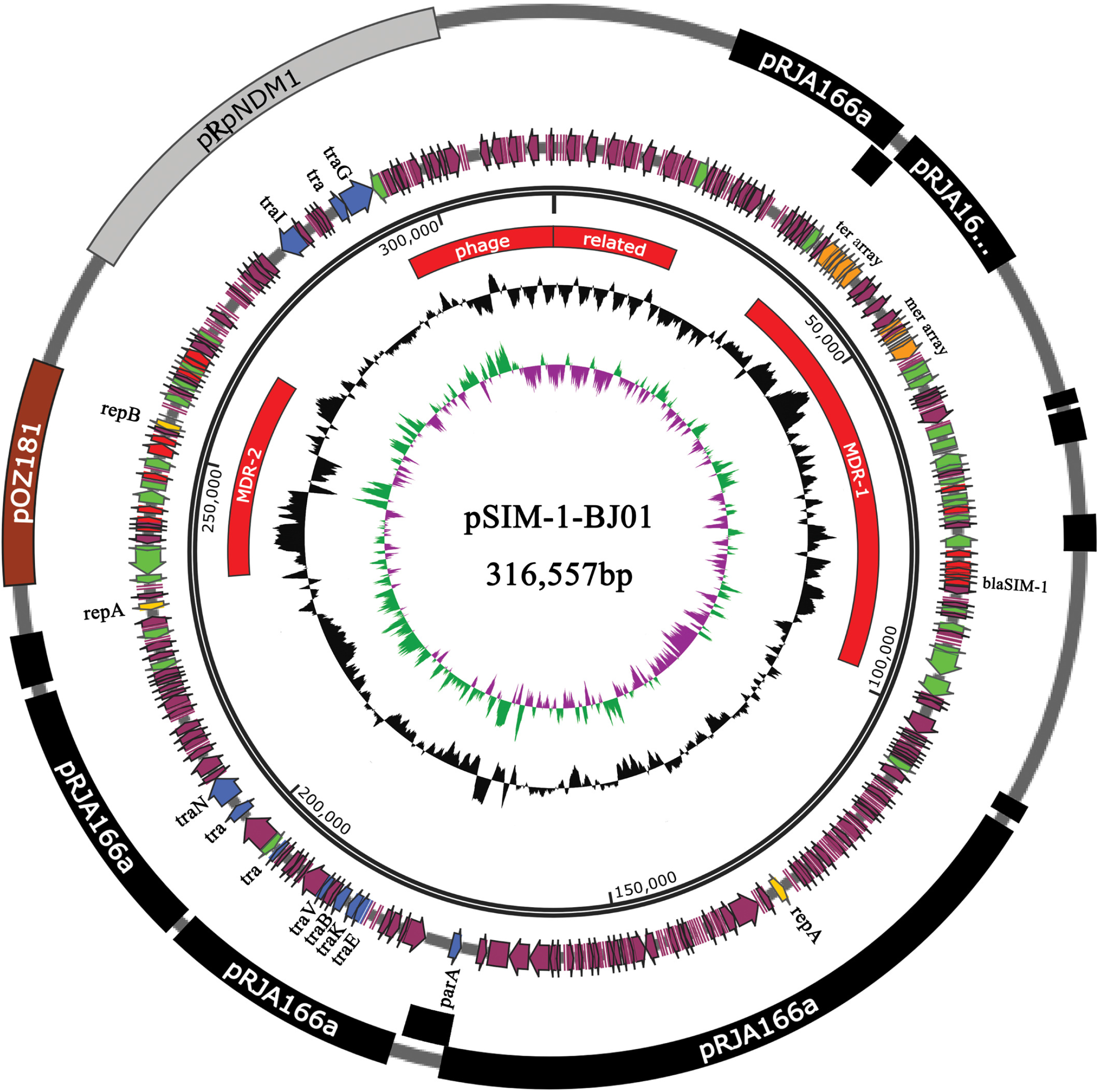
Schematic map of the plasmid pSIM-1-BJ01. The innermost circle presents GC-Skew [Green,GC skew+/Violet, GC skew -]. Starting from the inside, the second circle (black) presents GC content. The open reading frames (ORFs) are annotated in the fourth circle with arrows representing the direction of transcription and colored based on gene function classification [red, antibiotic resistance genes; yellow, replication initiation protein genes; blue, plasmid partition and conjugal transfer genes; green, transposon genes; orange, heavy-metal resistance genes]. The most outside circle indicates the backbone homologue of pSIM-1-BJ01 [black, the fragments from the plasmid pRJA166a; grey, the fragment from the plasmid pRpNDM1; brown, the fragment from the plasmid pOZ181].

**Figure 2.**
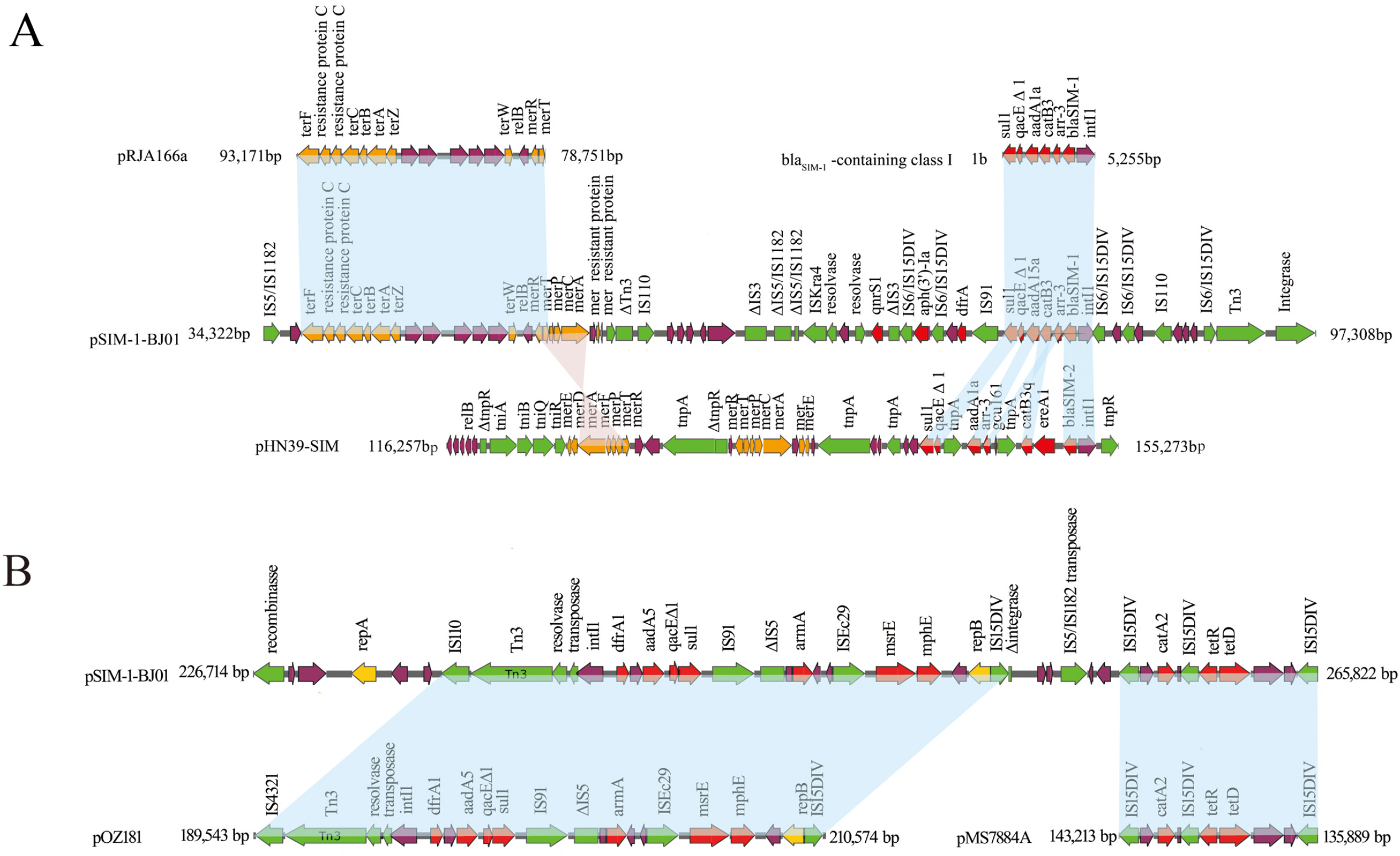
Organization and Alignment of the pSIM-1-BJ01 MDR regions. (A) The pSIM-1-BJ01 MDR-1 region. (B) The pSIM-1-BJ01 MDR-2 region. Genes are denoted by arrows and are colored based on gene function classification. Red, drug-resistant genes; green, mobile elements; orange, metal-related genes; violet, common genes.

**Figure 3.**
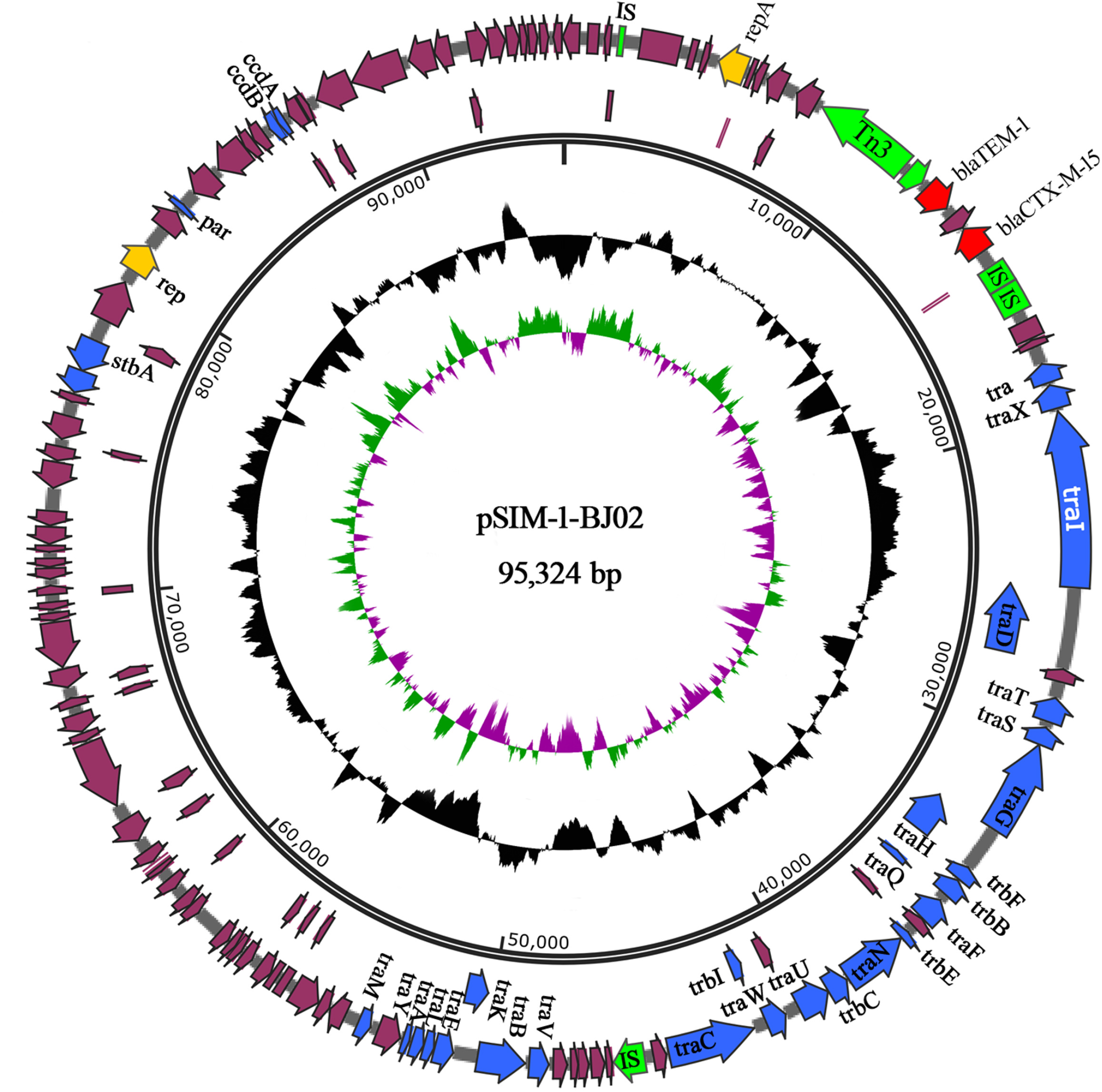
Schematic map of pSIM-1-BJ02. The innermost circle presents GC-Skew [Green,GC skew+/Violet, GC skew -]. Starting from the inside, the second circle (black) presents GC content. The open reading frames (ORFs) are annotated in the fourth circle with arrows representing the direction of transcription and colored based on gene function classification [red, antibiotic resistance genes; yellow, replication initiation protein genes; blue, plasmid partition and conjugal transfer genes; green, transposon genes].

**Figure 4.**
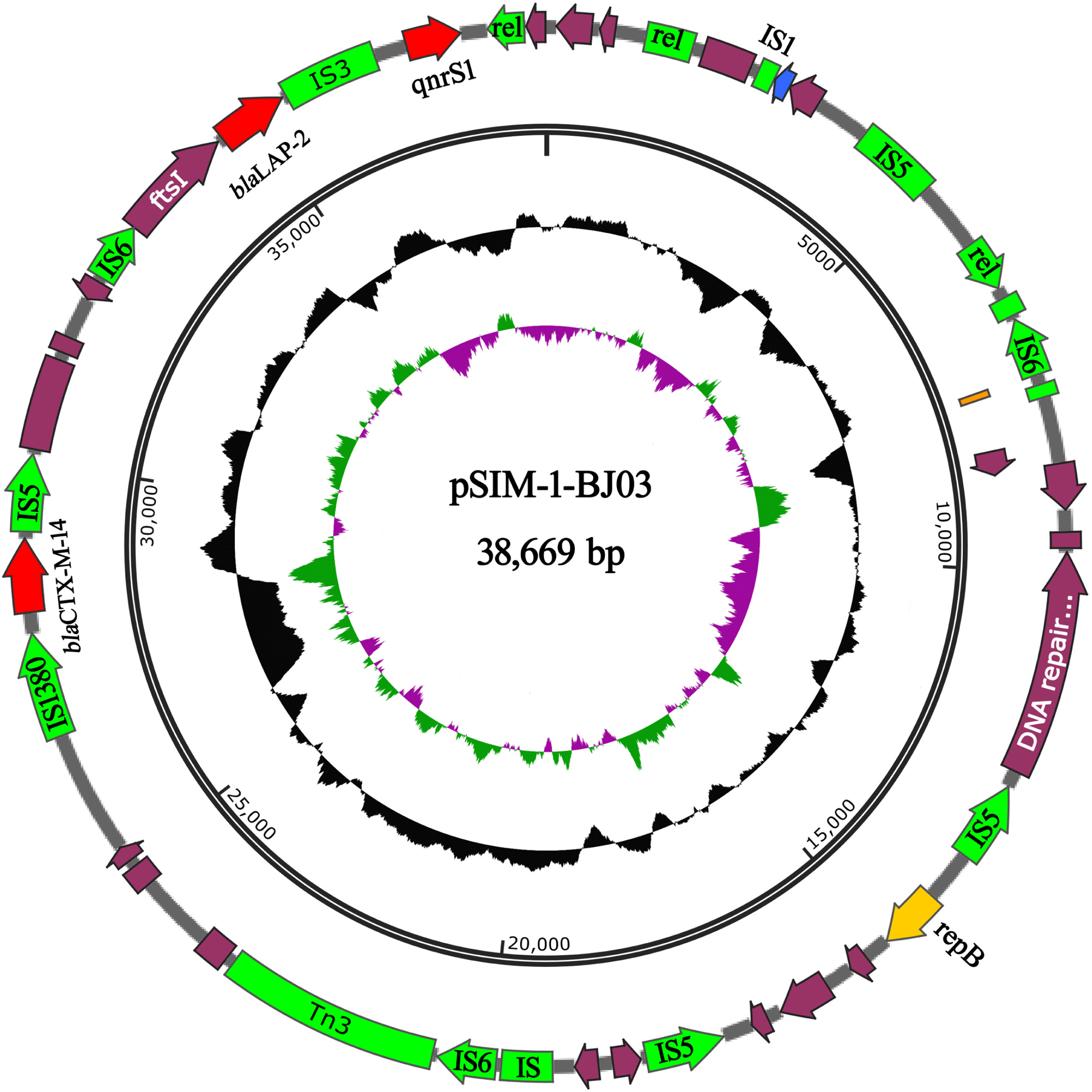
Schematic map of pSIM-1-BJ03. The innermost circle presents GC-Skew [Green,GC skew+/Violet, GC skew -]. Starting from the inside, the second circle (black) presents GC content. The open reading frames (ORFs) are annotated in the fourth circle with arrows representing the direction of transcription and colored based on gene function classification [red, antibiotic resistance genes; yellow, replication initiation protein genes; blue, plasmid partition and conjugal transfer genes; green, transposon genes].

### Overview and Comparative Analysis of pSIM-1-BJ01

The plasmid pSIM-1-BJ01 is a 316,557 bp circular plasmid containing 342 putative ORFs (153 hypothetical proteins), the average GC content is 46%. The new megaplasmid pSIM-1-BJ01 encodes three replication initiation proteins (two RepA and one RepB). In addition, it carries plasmid conjugal transfer proteins TraE, TraK, TraB, TraV, TraN, TraD, TraI, TraF and TraG, which code for the entire conjugation machinery. Two multidrug-resistance regions (MDRs) and one unique region related to phage invasion and assembly also existed in its backbone, as shown in Fig. 1.

The sequence alignment showed that most of the pSIM-1-BJ01 backbone was homologous to that of the plasmid pRJA166a (Genebank Acession Number CP019048.1) isolated from a clinical hypervirulent *K. pneumonia* ST23 strain at Shanghai of China recently (6). In addition, a small part of the backbone of pSIM-1-BJ01 is identical to that of the plasmid pOZ181 (Genebank Acession Number CP016764.1) isolated from *Citrobacter freundii* strain B38 at Guangzhou of China (7), and the other small part of the backbone of pSIM-1-BJ01 is identical to that of the plasmid pRpNDM1(Genebank Acession Number JX515588.1) isolated from *Raoultella planticola* strain RpNDM1 at Gansu of China (8), as shown in the outset circle of Fig. 1.

### The MDR-1 region of pSIM-1-BJ01

The plasmid pSIM-1-BJ01 possesses two MDR regions. The MDR-1 harboring the *bla*_SIM-1_ is totally 63-kb in length (from 34,322 bp to 97,308 bp), including nine resistant genes (*bla*_SIM-1_, *arr3, catB3, aadA15a, qacEΔ1, sulI, dfrA, qnrS1* and *aph(3’)-Ia*) as shown in Fig. 2A. The *bla*_SIM-1_ is also embedded in the conserved gene cassette which has 98% sequence identity to the reported class 1 integron from the earliest reported *A. baylyi* (Genebank Acession Number JF731030) or *A. baumannii* (Genebank Acession Number AY887066) (1), indicating a horizontal gene transfer of the conserved *bla*_SIM-1_ cassette from *Acinetobacter spp*. to *K. pneumonia*. The difference is that *aadA1* (streptomycin resistance gene) was replaced by *aadA15a* in the class 1 integron of pSIM-1-BJ01. AadA15a, displaying two amino acids substitution Met126Thr and Thr136Ile, is a new derivative of AadA15 (Genebank Acession Number WP_071846190.1) and has 93% amino acid sequence identity with AadA1. Except for the five resistant genes embedded in the class 1 integron, another three antibiotic resistant genes *dfrA*(trimethoprim resistant gene), *qnrS1* (quinolone resistance gene) and *aph(3’)-Ia* (kanamycin resistance gene) also existed in the MDR-1 region. The three resistant genes are respectively flanked by ISs at 5’ and 3’ ends, implying the active occurrence of the drug-resistant gene transfer.

Up to now, mentioning the *bla*_SIM_-harbouring plasmids, only the pHN39-SIM from *P. aeruginosa* was fully sequenced and it was untransferable to *E.coli* through conjugation or electroporation (4). It contains one MDR region with seven resistant genes (*bla*_SIM-2_, *ereA1, catB3q, arr3, aadA1a, qacEΔ1* and *sulI*) and two different mercury resistance gene (mer) arrays (a and b). Compared with the pHN39-SIM, the MDR-1 region of pSIM-1-BJ01 owns one *mer* resistance gene array. The *mer* array of pSIM-1-BJ01 has 78% sequence similarity to the *mer* array b of pHN39-SIM, as shown in Fig. 2A. Except for the *mer* array, the pSIM-1-BJ01 also owns the tellurium resistant gene (*ter*) array in MDR-1 region, the *ter* array of pSIM-1-BJ01 is 99% identical to that of pRJA166a (from 78,751bp to 93,171bp) (6).

The *relB* adjacent to the *mer* array belongs to type II toxin-antitoxin systems which are expected to increase the longevity of plasmids in bacterial populations even in the absence of selection pressure due to antibiotics and metals. Co-occurrence of metal resistance genes, antibiotic resistance genes and toxin-antitoxin genes together on a conjugative plasmid highlights the potential clinical consequences of co-selection (9).

### The MDR-2 region of pSIM-1-BJ01

The length of MDR-2 is 39-kb from 226,714 bp to 265,885 bp, including ten resistant genes (*msrE, mphE, armA, dfrA1, aadA5, qacEΔ1, sul1, catA2, tetR* and *tetD*) as shown in Fig. 2B. The MDR-2 contains two replication initiation protein genes *repA* and *repB*. One fragment of the MDR-2 (from 233,359 bp to 254,517 bp) is 99% identical to that of the plasmid pOZ181(from 189,543 bp to 210,753 bp) isolated from a *bla*_IMP-4_-carrying *C.freundii* B38 (7), the 21-kb fragment contains resistant genes *msrE* (the macrolide efflux pump gene), *mphE* (the macrolide 2’-phosphotransferase gene), *armA* (16s rRNA methylase gene) and sul1-type class I integron which has the *dfrA1-gcu37-aadA5-qacEΔ1-sul1* cassette. The only difference between the two drug-resistant fragments is that the *IS4321* at the upstream of sul1-type integron on the pOZ181 is replaced by *IS110* at the corresponding part on pSIM-1-BJ01. The other fragment (from 258,498 bp to 265,885 bp), carrying resistant genes *catA2, tetR* and *tetD*, is 100% identical to that (from 143,213 bp to 135,889 bp) of the *bla*_IMP-4_-carrying pMS7884A (Genebank Acession Number CP022533.1) isolated from an *Enterobacter cloacae*.

### Overview of pSIM-1-BJ02

pSIM-1-BJ02 is a 95,324 bp circular plasmid containing 124 putative ORFs (54 hypothetical proteins), and the average GC content is 52%, as shown in Fig. 3. pSIM-1-BJ02 carrying *bla*_TEM-1_ and *bla*_CTX-M-15_ is not a new plasmid, and it is 100% homologous to the plasmid pL22-5 (Genebank Acession Number CP031262.1) isolated from a hypermucoviscous strain *Klebsiella quasipneumoniae* ST367 in China and is 99% identical to the plasmid pKF3-94 (Genebank Acession Number FJ876826) isolated from a clinical drug-resistant strain *K. pneumonia* KF3 in China (10).

### Overview of pSIM-1-BJ03

The plasmid pSIM-1-BJ03 is a 38,669 bp circular plasmid containing 29 putative ORFs (8 hypothetical proteins), and the average GC content is 49%, as shown in Fig. 4. The small pSIM-1-BJ03 carrying *bla*_LAP-2_ and *bla*_CTX-M-14_ is firstly reported here, its multidrug-resistant region (from 25,868 bp to 38,669 bp) is 100% identical to that of pE66An (Genebank Acession Number HF545433.1) (from 80105 bp to 67304 bp) isolated from *E.coli* in Vietnam.

## DISCUSSION

The infection rate of carbapenem-resistant *K. pneumoniae* (CRKP) has increased substantially in the past 10 years, even a fatal outbreak of hypervirulent carbapenem-resistant *K. pneumonia* (hv CRKP) in a Chinese hospital (11-13). Here, we presented a clinically isolated *K. pneumonia* with three resistance plasmids. Among them, the plasmid pSIM-1-BJ01 carried the rare carbapenemase gene *bla*_SIM-1_.

The very rare carbapenem-resistant gene *bla*_SIM-1_ was usually found in *Acinetobacter spp*. and *Pseudomonas aeruginosa* isolated from South Korea and China, and the *bla*_SIM-1_-carrrying megaplasmid pSIM-1-BJ01 isolated from *K. pneumonia* was less reported. Most of the pSIM-1-BJ01 backbone were homologus to the corresponding parts of pRJA166a, which was isolated from a clinical hypervirulent *K. pneumonia* ST23 strain at Shanghai of China recently. The plasmid pRJA166a carrying *bla*_DHA-1_ was able to disseminate across an expanded collection of *K. pneumonia* and keep its stability across the transconjugants. The highly homologous backbones between the two plasmids imply the close relationship of evolution and reflect the active happening of gene convergence under clinical selection pressure.

Unlike the reported *bla*_SIM_-harbouring plasmid pHN39-SIM from *P. aeruginosa*, the plasmid pSIM-1-BJ01 was able to transfer to *E.coli*, indicating the possible spread of *bla*_SIM_ among Enterobacteriaceae. Now there was one case report about the *bla*_SIM-1_ carrying *E.coli* from India (14). The emergency of the multidrug-resistant megaplasmid pSIM-1-BJ01 reflects the active occurrence of resistant gene convergence under clinical selection pressure, we should pay more attention to supervise the dissemination of *bla*_SIM_ in Enterobacteriaceae.

Except for the new megaplasmid pSIM-1-BJ01, the moderate-size plasmid pSIM-1-BJ02 is a common reported plasmid isolated from clinical hypermucoviscous drug-resistant *Klebsiella spp*. in Chinese hospitals; the small-size plasmid pSIM-1-BJ03 was first reported here with its MDR region identical to that of pE66An isolated from *E.coli* in Vietnam; so it is urgent to supervise these resistant plasmids’ transmission and evaluate the epidemicity among clinical *Klebsiella spp*. and Enterobacteriaceae.

## AUTHOR CONTRIBUTIONS

HHy and WSw: conception and design of the study. LY, LH, ZW and LJ: acquisition of data. LY: genome sequence analysis and interpretation of data, drafting the article. LH, ZW AND LJ: sample isolation and identification, antimicrobial resistance analysis and molecular analysis.

## Transparency declarations

None to declare.

